# Evaluation of Machine Learning predictions of a highly resolved time series of Chlorophyll-a concentration

**DOI:** 10.1101/2021.05.12.443749

**Authors:** Felipe de L. L. de Amorim, Johannes Rick, Gerrit Lohmann, Karen Helen Wiltshire

## Abstract

Pelagic Chlorophyll-a concentrations are key for evaluation of the environmental status and productivity of marine systems. In this study, chlorophyll-a concentrations for the Helgoland Roads Time Series were modeled using a number of measured water and environmental parameters. We chose three common Machine Learning algorithms from the literature: Support Vector Machine Regressor, Neural Networks Multi-layer Perceptron Regressor and Random Forest Regressor. Results showed that Support Vector Machine Regressor slightly outperformed other models. The evaluation with a test dataset and verification with an independent validation dataset for chlorophyll-a concentrations showed a good generalization capacity, evaluated by the root mean squared errors of less than 1 μg L^−1^. Feature selection and engineering are important and improved the models significantly, as measured in performance, improving by a minimum of 48% the adjusted R^2^. We tested SARIMA in comparison and found that the univariate nature of SARIMA does not allow for better results than the Machine Learning models. Additionally, the computer processing time needed was much higher (prohibitive) for SARIMA.

## Introduction

Pelagic Chlorophyll-a concentrations (chl-a) are a common indicator of primary production and key to evaluation of the health and productivity of marine and freshwater systems [1],[2]. It is therefore of crucial importance to accurately measure/ predict chlorophyll from proxy parameters in such systems [3]. Accelerated global warming is exacerbating climate change and unsettling ecosystems processes, while the impacts directly affect the marine primary production triggering an upwards transfer of effects which reach humans. Thus, the importance of modelling model chlorophyll is emphasized in environments undergoing change resulting from global warming [4].

Prediction of chlorophyll-a time series data is a challenge due their complexity and nonlinearity, and indeed, conventional approaches show limitations with prediction of unobserved data [5],[6]. To date, all conventional approaches including factors based on single measurements are limited with regard to prediction accuracy of Chlorophyll-a concentrations [7]. A few previous studies have tried to implement various machine learning techniques to predict chlorophyll concentrations, mainly in fresh water systems, with a few in marine regions [8],[9],[10],[11].

Machine Learning (ML) techniques constitute a set of tools belonging to the field of Computer Science and Artificial Intelligence. The versatility of these techniques allow the successful application in many fields of science and to a great variety of problems. The focus is often placed on tackling pattern recognition problems and on the construction of predictive models to make data-driven decisions [12]. According to [13], the general benefits of ML algorithms for time series prediction over classical methods include the ability of supporting noisy features, noise and complexity in the relationships between variables and in the handling of irrelevant features.

State-of-the-art ML algorithms for time series regression include Random Forest Regressor (RF), Support Vector Machine Regressor (SVR) and Neural Networks Multi-layer Perceptron Regressor (MLP). All of these have been used to some degree in the literature for the prediction of chlorophyll-a concentrations in aquatic systems, and achieve significantly accurate results in both error and goodness of fit metrics [3],[11],[14]. These are studies based in chl-a time series either with short length and daily frequency or long term low frequency sampling time series, using different ML methods to best predict chl-a behavior. The features applied as predictors in these studies are limited to just a few, but it must be considered that the dynamics in lacustrine systems are distinct from the ones presented in marine systems. Here we extend these ideas and test these methods on a good quality long-term time series, the Helgoland Roads Time Series, evaluating the prediction using unseen data. With the purpose to compare ML methods with a classical statistical regression model, we included an improved Autoregressive Integrated Moving Average (ARIMA) model, called Seasonal ARIMA (SARIMA), which includes seasonal parameters to support data with a seasonal component [15].

The objective of this work is to evaluate the accuracy of Machine Learning algorithms for the estimation of Chlorophyll-a concentration, using in situ high resolution long term datasets. We (1) assess three ML algorithms – Random Forest, Support Vector Regressor and Neural Networks Multi-layer Perceptron Regressor – for Chlorophyll-a concentration estimation; (2) examine the importance of feature selection and engineering in the different models and (3) compare with and evaluate a univariate SARIMA classical regression model.

## Methods

All the ML models used in this study were implemented applying the ‟Scikit-Learn package”, which is an open-source Python module project that integrates a wide range of common ML algorithms [16],[17], while the SARIMA model was implemented with the statsmodels package [18]. The pre-processing was also implemented in the Python environment, using the well-known packages Pandas, Numpy and Scipy [19].

### Datasets

The Helgoland Roads is a long term pelagic monitoring site (54°11.3’ N, 7°54.0’ E) about 60 km off the German coast and represents a marine transition zone between coastal waters and open sea (Fig 1) [20]. Since 1962, surface water samples are collected on working days, taken with a bucket lowered from a research vessel. Secchi depth and water temperature (SST) are measured in situ and the water samples analyzed in the laboratory for nutrients (nitrate, phosphate and silicate), and salinity. Chlorophyll-a concentration measurements started in the end of 2001, acquired in laboratory by FluoroProbe (bbe Moldaenke GmbH, Kiel, Germany) [21] and since 2004 have been complemented with High Performance Liquid Chromatography analysis (HPLC) [22],[23].

**Fig 1.**
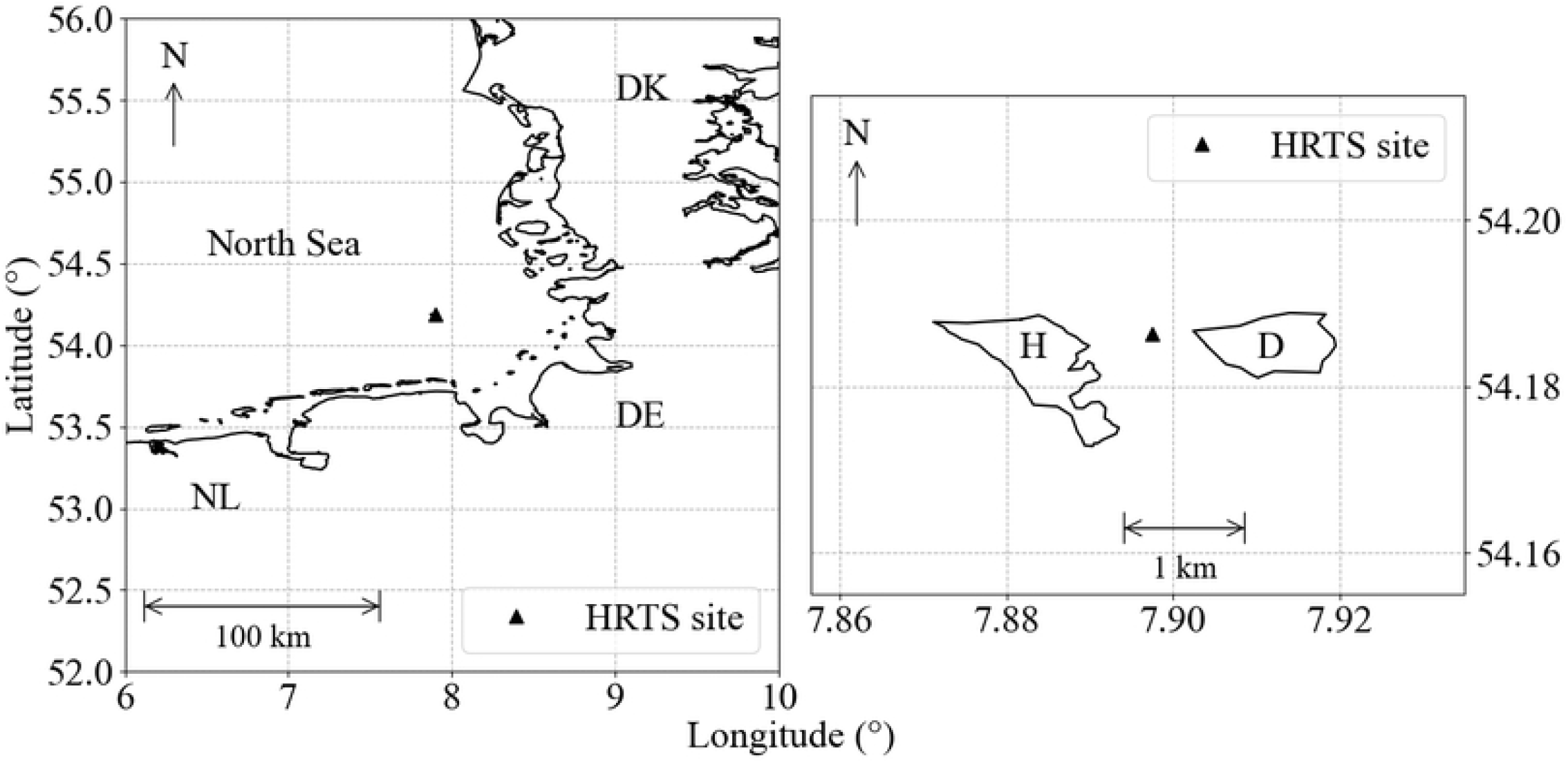
Helgoland Roads monitoring site position (black triangle) in the German Bight, between the Helgoland (H) and Dune (D) islands.

Sunshine duration, wind speed and direction [24],[25],[26], North Atlantic Oscillation (NAO) daily index (NOAA ESRL Physical Sciences Laboratory, Boulder, Colorado, USA, 2020) and zooplankton abundance [27], were added to the Helgoland Roads parameter matrix for this work (Table 1). As indicated in the literature [28],[29],[30] and also from working experience, the included parameters are environmental variables which determine algal verdure and thus modulate chlorophyll-a concentrations in marine systems.

**Table 1.** Statistical description of parameters used as determinants to predict chlorophyll-a concentration (target) after linear interpolation (std, min and max are standard deviation, minimum and maximum values) respectively.

| Parameters | Units | Counts | Mean | std | min | max | Median |
| --- | --- | --- | --- | --- | --- | --- | --- |
| Secchi depth | m | 4920 | 3.70 | 1.80 | 0.20 | 12.00 | 3.67 |
| SST | °C | 4920 | 10.64 | 5.00 | 1.10 | 20.00 | 10.30 |
| Salinity | — | 4920 | 32.31 | 1.06 | 26.71 | 36.11 | 32.42 |
| SiO<sub>4</sub> | μmol L<sup>-1</sup> | 4920 | 6.49 | 5.04 | 0.01 | 37.20 | 5.26 |
| PO<sub>4</sub> | μmol L<sup>-1</sup> | 4920 | 0.56 | 0.41 | 0.01 | 3.98 | 0.53 |
| NO<sub>3</sub> | μmol L<sup>-1</sup> | 4920 | 10.19 | 9.82 | 0.10 | 77.38 | 7.12 |
| Sunlight Duration | h | 4920 | 4.78 | 4.52 | 0.00 | 16.60 | 3.60 |
| NAO index | — | 4920 | -9.44 | 121.55 | -564.29 | 351.95 | -0.52 |
| Wind Direction | Degrees [°] | 4920 | 203.09 | 74.71 | 20.00 | 353.00 | 212.00 |
| Wind Speed | m s<sup>-1</sup> | 4920 | 8.23 | 3.28 | 1.60 | 20.80 | 7.80 |
| Zooplankton Abundance | individuals m<sup>-3</sup> | 4920 | 3164.70 | 4450.64 | 5.00 | 75364.50 | 1676.21 |
| Chlorophyll-a | µg m<sup>-1</sup> | 4920 | 2.40 | 2.86 | 0.00 | 45.45 | 1.48 |

### Data Pre-processing

The raw data of Helgoland Roads is characterized by long term measurements on work-daily frequency, with missing values during weekends and extreme bad weather days. When merged by date with other features like zooplankton abundance, it ends with approximately 40% of missing data in the time series. To fill the missing data, creating a regular sampled daily time-series, a number of imputation methods were tested in sunlight duration, a feature added to the Helgoland Roads from an external source, with no missing values. After creating a synthetic missing values dataset with sunlight duration, we calculated root mean square error (RMSE) and coefficient of determination (R^2^) between the original and interpolated data. Minimum changes in frequency distribution between missing data and interpolated variable, lowest RMSE and highest R^2^ based the decision to use a Linear Interpolation, supported by [30]. After the interpolation, we have a daily dataset comprising approximately 13 years, from 02/11/2001 to 22/04/2015.

Additionally, date features were generated namely ‟year” and ‟day of year” from 1 to 365 or 366. The cyclic variables: day of year and wind direction were transformed with (sin, cos) [2π (day, degrees) / (number of days in year, 360)] to ensure that the last day of a year is understood to be in sequence with the first day of the next year and 0° degree in direction is equal to 360° [31].

In this study, to validate the performance of the ML models, the dataset was split in 80% (n = 3940) for model training, and 20% (n = 980) for model testing, so we could investigate the model generalization ability [32]. To eliminate the dimensional differences of the data and also improve the prediction ability of the models, we used the StandardScaler method from the Scikit-Learn package, which standardizes features by removing the mean and scaling to unit variance.

The training dataset, the sample of data used to fit the model, dates from 02/11/2001 to 15/08/2012 (~11 years), while the test set is from 16/08/2012 to 22/04/2015 (~2.5 years) and it is used for model evaluation (Fig 2). For independent validation, we used a linear interpolated time series of HPLC estimated chlorophyll data (05/05/2015 to 27/11/2018, n=348).

**Fig 2.**
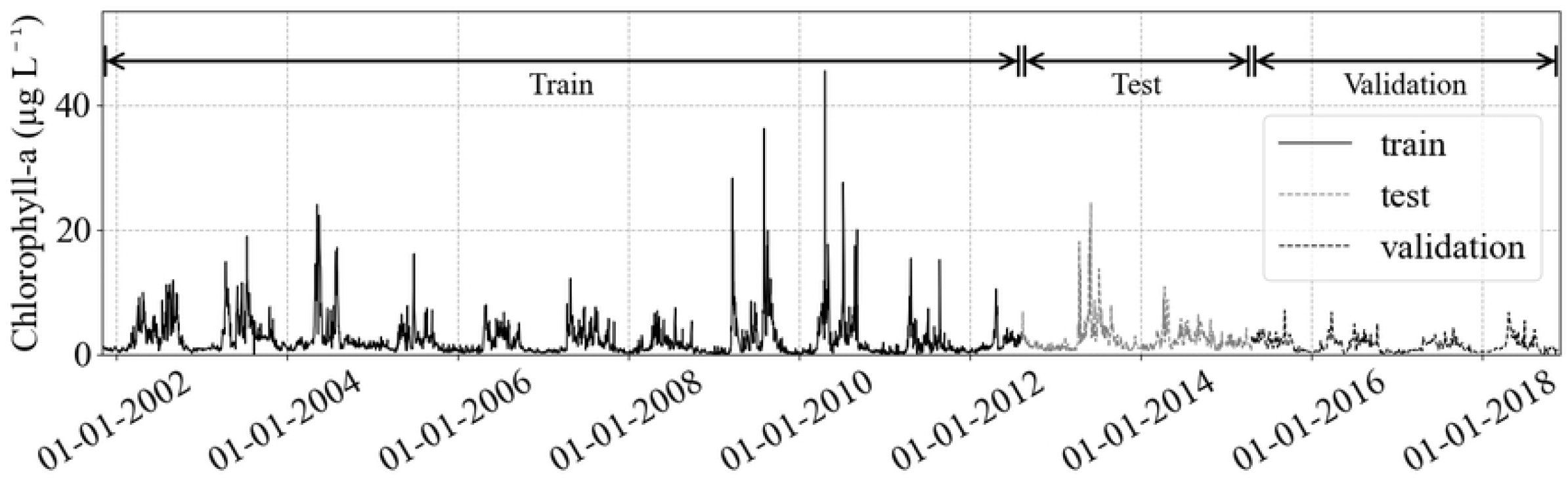
The train and test partition in chlorophyll-a concentration target (black solid and gray solid lines, respectively), and the HPLC chl-a validation dataset (black dashed). After the split, the testing dataset will remain untouched, to guarantee no leakage of information to the training step. The validation dataset is the independent validation.

### Feature Engineering and Selection

The Pearson correlation coefficients were calculated to investigate linear relationships between Chlorophyll-a concentration and the other variables (Table 2). All correlation coefficients were lower than 0.5, indicating no strong linear correlation between chlorophyll and any other variable.

**Table 2.** Pearson correlation among predictors and the target chlorophyll-a concentration.

| Predictors | Code | Correlation |
| --- | --- | --- |
| year | year | -0.05 |
| sin(days) | sin(days) | 0.04 |
| cos(days) | cos(days) | -0.46 |
| Secchi depth | SD | 0.15 |
| Sea Surface Temperature | SST | 0.27 |
| Salinity | Salinity | -0.22 |
| Silicate | SiO4 | -0.31 |
| Phosphate | PO4 | -0.29 |
| Nitrate | NO3 | -0.09 |
| Sunlight duration | Sunlight | 0.31 |
| NAO index | NAO | 0.06 |
| sin(wind direction) | sin | 0.02 |
| cos(wind direction) | cos | 0.10 |
| Wind Speed | Speed | -0.20 |
| Zooplankton Abundance | Abundance | 0.22 |
| Chlorophyll-a | Chl | 1.00 |

Prediction is a major task of time series data mining, which uses known historical values to estimate future values, and feature selection and engineering is essential and crucial for accurate predictions [33]. To seek improvement, 15 days lagged predictors were generated, totalizing 211 features [34]. The choice of lags was based in a two-week period where all the predictors supposedly influence chlorophyll-a concentration, including chl-a lagged values, i.e., the lagged target values were used as predictors. As there are significant seasonal differences e.g., summer and winter nutrients uptake, the definition of two weeks seemed reasonable for this work to input information, considering that the Machine Learning algorithms are data-driven and they are not mechanistic models [35].

A large number of features in the dataset drastically affects both the training time as well as the accuracy of machine learning models. One means to limit model complexity from multiple variables is to reduce the model by selectively eliminating predictors. Feature selection procedure was conducted applying a simple sequential addition of lagged predictors, and Recursive Feature Elimination. For the latter, we use Scikit Learn module Recursive Feature Elimination with cross validation (Scikit-Learn feature.selection RFECV module) and Ridge estimator, to estimate the best number of features balanced with accuracy (Fig 3). After the best number of features were defined with the Ridge cross-validation method, we applied Recursive Feature Elimination (Scikit-Learn feature.selection RFE module) with SVR linear estimators, this way selecting the 17 best parameters to model chl-a in a robust manner [36].

**Fig 3.**
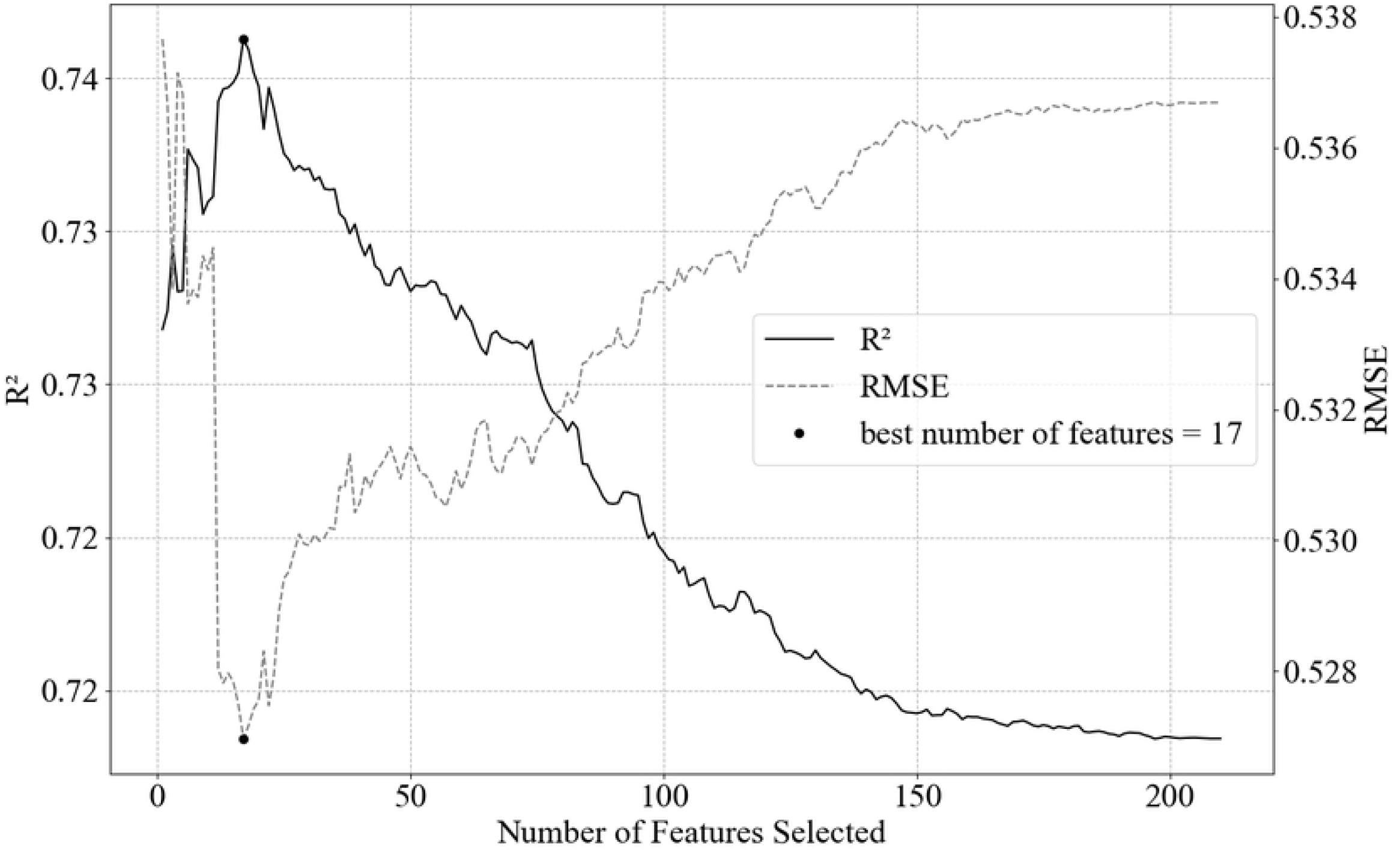
Result of RFECV with Ridge estimator. The black dot represents the maximum value of 17 selected features (predictors) to reach the highest explained variance. After the maximum value, there is an exponential decay/increase in the R^2^/RMSE.

### Model selection and parameter tuning

The algorithms evaluated in this study are Random Forest Regressor (RF) [37], Support Vector Machine Regressor (SVR) [38] and Multi-layer Perceptron Regressor Neutral Network (MLP) [39],[40]. Depending upon the study cases, different ML algorithms usually require some adjustments. These are often crucial for the development of a successful application. Each ML algorithm has parameters so-called hyperparameters, which define the setup of the machine to modelling the target function. For each model, a search range of hyperparameters was tested. In case where a value was selected at the edge of the search range, a new cross-validation was conducted including more values.

All hyperparameter tuning of the models (Table 3) is based on GridSearchCV in the Scikit-Learn package, which can evaluate all possible given combinations of hyperparameter values using 10-fold cross-validation to determine the best combination of hyperparameter that has the best accuracy of the model in terms of coefficient of determination (R^2^). Cross-validation is model validation techniques for obtaining reliable and stable models. The use of multiple models in the evaluation removes possible biases of some models with some data sets. We use the training dataset to search for the best parameters, and report the prediction performances on the test dataset using these parameters [41]. The mentioned grid search was performed independently for each model on the training subset.

**Table 3.** Hyperparameter tested in GridSearchCV and the ones applied to each ML algorithms.

| Model | Hyperparameter | Selected value | Default |
| --- | --- | --- | --- |
| SVR | kernel | rbf | rbf |
|  | C | 3 | 1 |
|  | gamma | 0.01 | 0.1 |
|  | Epsilon | 0.1 | 0.0001 |
| MLP | max_iter | 90 | 200 |
|  | hidden_layer_sizes | 40 | 100 |
|  | activation | logistic | relu |
|  | solver | adam | adam |
|  | Alpha | 0.5 | 0.0001 |
|  | warm_start | True | False |
| RF | bootstrap | True | True |
|  | max_depth | 25 | None |
|  | max_features | 21 | auto |
|  | min_samples_leaf | 9 | 1 |
|  | min_samples_split | 2 | 2 |
|  | n_estimators | 80 | 100 |

R^2^, adjusted coefficient of determination (adj R^2^) and RMSE are the metrics that were used in this work to evaluate the predictions. The use of adj R^2^ in multiple regression is important because it increases only when new independent variables that increase the explanatory power of the regression equation are added; making it a useful measure of how well a multiple regression equation fits the sample data. A linear base model, available in Scikit-Learn module as ‟linear_model”, was used to observe the improvements using the more sophisticated algorithms.

### SARIMA Model

For the SARIMA model, the univariate chl-a data was used, while maintaining the partitions in the training and test dataset. To test stationarity, the Augmented Dickey-Fuller test (ADF) was applied indicating significant stationarity (p<0.05) in the train and test datasets. To fill the model (p,d,q)x(P,D,Q)_365_, where 365 represents the seasonality, the best auto-regressive (p, P) and moving average (q, Q) parameters were selected using an iterative method in the train dataset. The parameters ranged from 0 to 4 in the non-seasonal parameters (p,q) and 0 to 2 in the seasonal parameters (P, Q), selecting the combination with lowest Akaike Information Criteriation (AIC). The difference order parameters d and D were 0, due to the stationarity results of the ADF test. The best parameters selected using the training dataset were (4, 0, 1) x (2, 0, 1)_365_, and this SARIMA model was used to fit the test dataset.

## Results

For this study, the best R^2^, adj R^2^ and RMSE achieved predicting chlorophyll-a using Support Vector Machine Regressor, Random Forest Regressor, and Neural Network Multi-layer Perceptron Regressor are presented in Table 4. In a combination of hyperparameters tuning and feature selection, SVR reached the best R^2^ (0.78) and RMSE (1.113 μg L^−1^) compared with the other algorithms. However, these were slightly better results. The algorithms presented good performances for the training data set during the cross-validation step (Fig 4). In addition, the predicted values were close to the observed data (Fig 5).

**Table 4.** Comparison of non-optimized (Default) and optimized model performances for predicting Chlorophyll-a concentration during training (train) and testing (test) steps. The linear model serves as a base model.

| Default |  |  |  |  |  |  |
| --- | --- | --- | --- | --- | --- | --- |
|  | train |  |  | test |  |  |
|  | adj R<sup>2</sup> | R<sup>2</sup> | RMSE (µg L<sup>-1</sup>) | adj R<sup>2</sup> | R<sup>2</sup> | RMSE (µg L<sup>-1</sup>) |
| SVM | 0.81 | 0.82 | 1.255 | 0.63 | 0.71 | 1.273 |
| RF | 0.96 | 0.96 | 0.607 | 0.64 | 0.71 | 1.258 |
| NN | 1 | 1 | 0.128 | 0.14 | 0.33 | 1.934 |
| Optimized |  |  |  |  |  |  |
|  | train |  |  | test |  |  |
|  | adj R<sup>2</sup> | R<sup>2</sup> | RMSE(µg L<sup>-1</sup>) | adj R<sup>2</sup> | R<sup>2</sup> | RMSE (µg L<sup>-1</sup>) |
| SVM | 0.77 | 0.77 | 1.424 | 0.77 | 0.78 | 1.113 |
| RF | 0.84 | 0.84 | 1.18 | 0.74 | 0.75 | 1.167 |
| NN | 0.75 | 0.75 | 1.474 | 0.76 | 0.76 | 1.144 |
| Linear (base model) |  |  |  |  |  |  |
|  | train |  |  | test |  |  |
|  | adj R<sup>2</sup> | R<sup>2</sup> | RMSE (µg L<sup>-1</sup>) | adj R<sup>2</sup> | R<sup>2</sup> | RMSE (µg L<sup>-1</sup>) |
|  | 0.74 | 0.76 | 1.47 | 0.65 | 0.73 | 1.227 |

**Fig 4.**
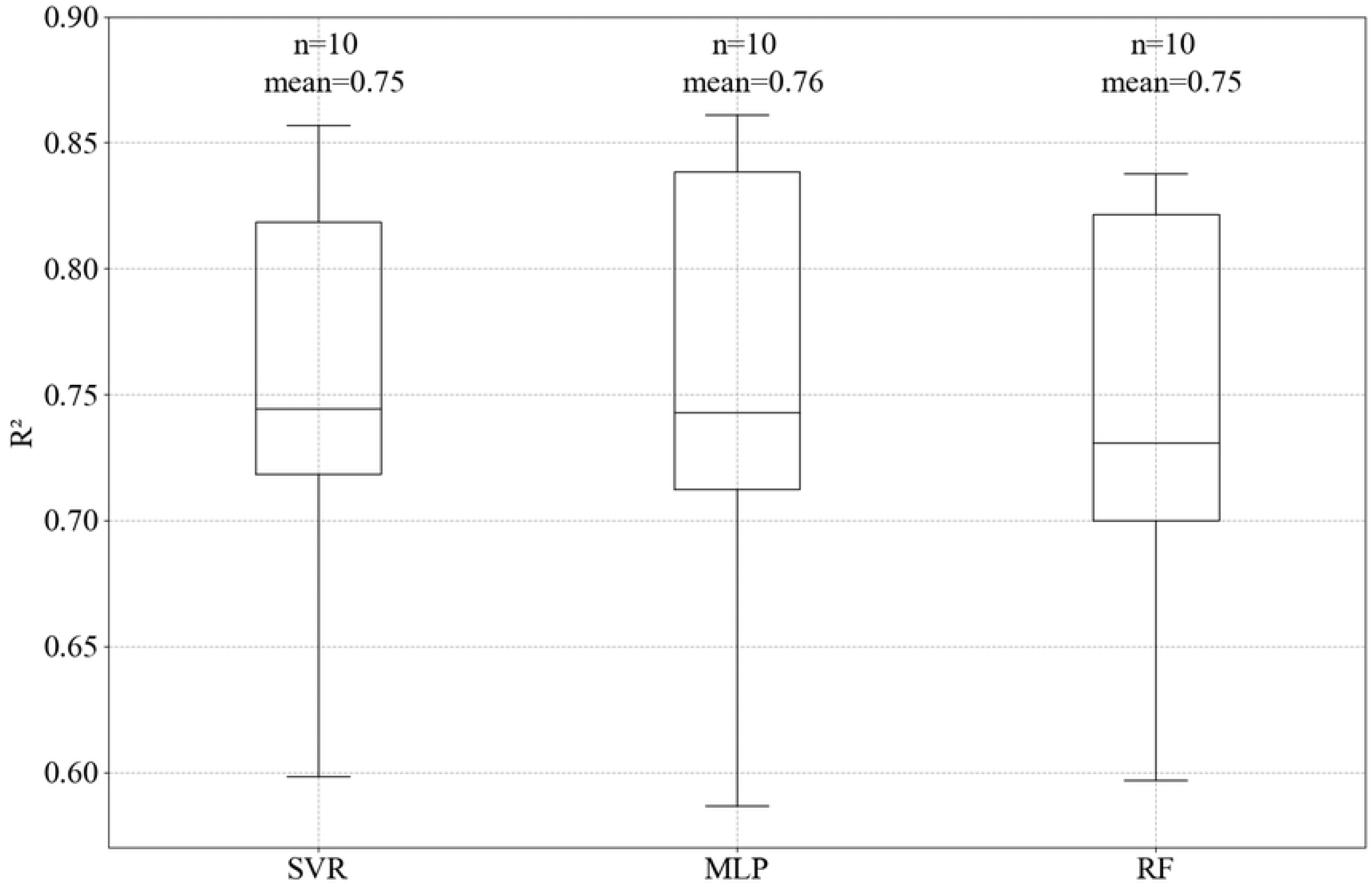
Boxplot of accuracy in the 10 fold cross-validation training step for the SVR, MLP and RF models, showing the mean and the number of folds (n) or subsets in the training data used to define the best hyperparameters.

**Fig 5.**
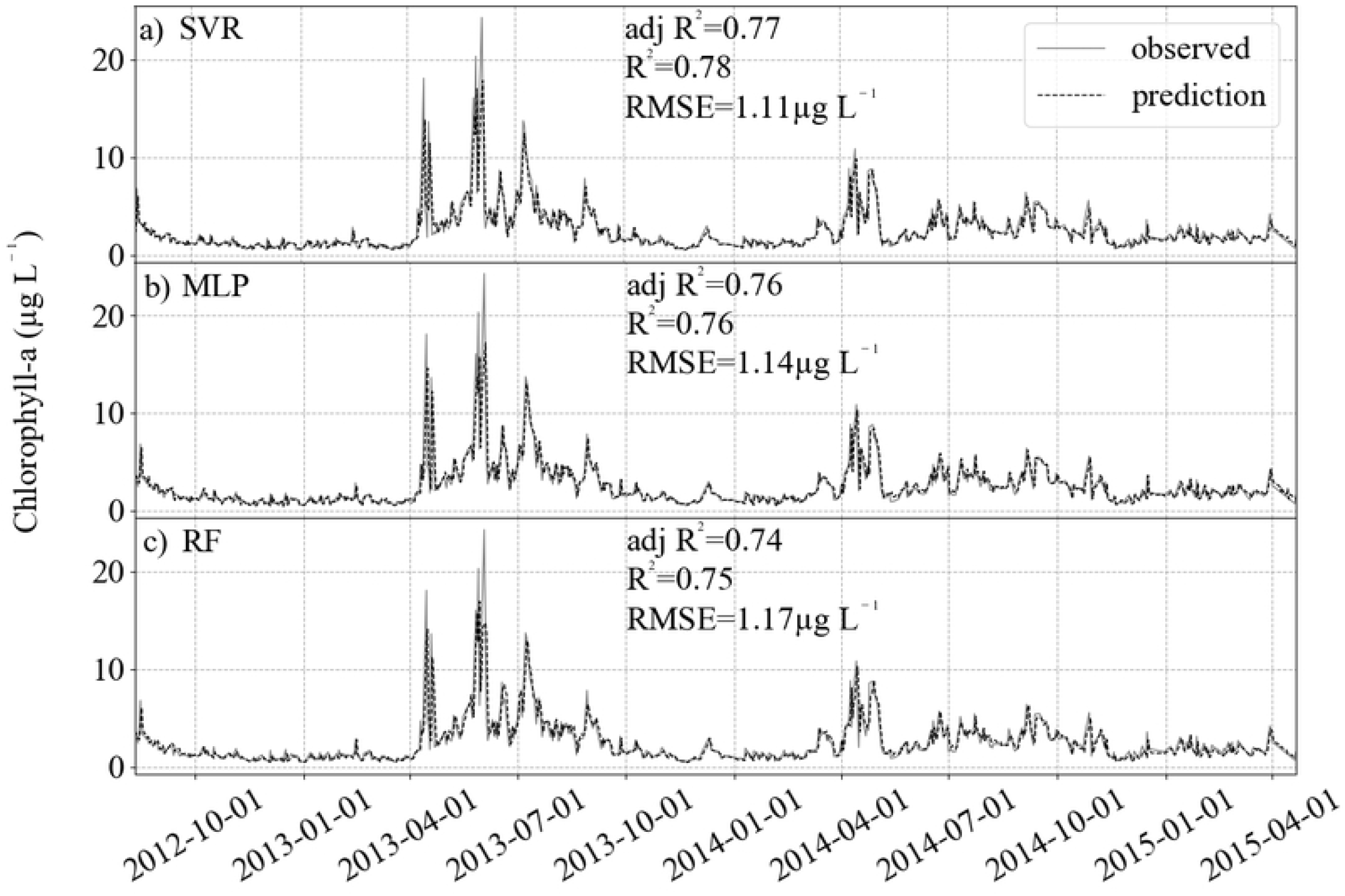
Result of prediction (black dashed) and comparison with the test dataset (gray solid). For the three algorithms, R^2^ is higher than 0.7 and RMSE lower than 1.2 μg L^−1^. a) SVR, b) MLP and c) RF.

The algorithms gave a good performance for the training dataset and allowed a good generalization for the test dataset, as can be seen from how close the predicted values are from the observed ones in Fig 5. Using all the 211 features and the default hyperparameters, the results in the test data were not as good as those from the optimized models (Table 4).

Considering the features used as inputs in each of the algorithms: for Random Forest the best result was acquired with the original features plus 2 lags, i.e., two past data in the time series, while with MLP the original features and 1 lag was enough to get the best achievable results for this algorithm in this study. For SVR, the Recursive Feature Elimination was implemented by combining Ridge and SVR linear estimators and selecting a maximum number of 17 predictors. This generated the following result: [‘SD, ‘SST’, ‘Salinity’, ‘SD_-1’, ‘SST_-1’, ‘SST_-2’, ‘SST_-9’, ‘SST_-12’, ‘SST_-13’, ‘SST_-14’, ‘SST_-15’, ‘Salinity_-1’, ‘Chl_-1’, ‘Chl_-4’, ‘Chl_-5’, ‘Chl_-7’, ‘Chl_-8’], with the negative numbers in the codes (Table 2) representing the applied lag in days. The adj R^2^ results, which are sensitive to the number of used predictors, show improvement from 0.14 to 0.76 for MLP, while for SVR was 0.63 to 0.77 and 0.64 to 0.74 to RF.

For the independent validation, a chl-a dataset acquired by HPLC, the predictions had better RMSE and R^2^ than the test datasets (Fig 6). Again, the higher values had limitations in prediction, but the lower variance compared with the training and testing datasets allowed better evaluation indicators, with RMSE for all algorithms in the order of 0.3 μg L^−1^ and R^2^ reaching approximately 0.9.

**Fig 6.**
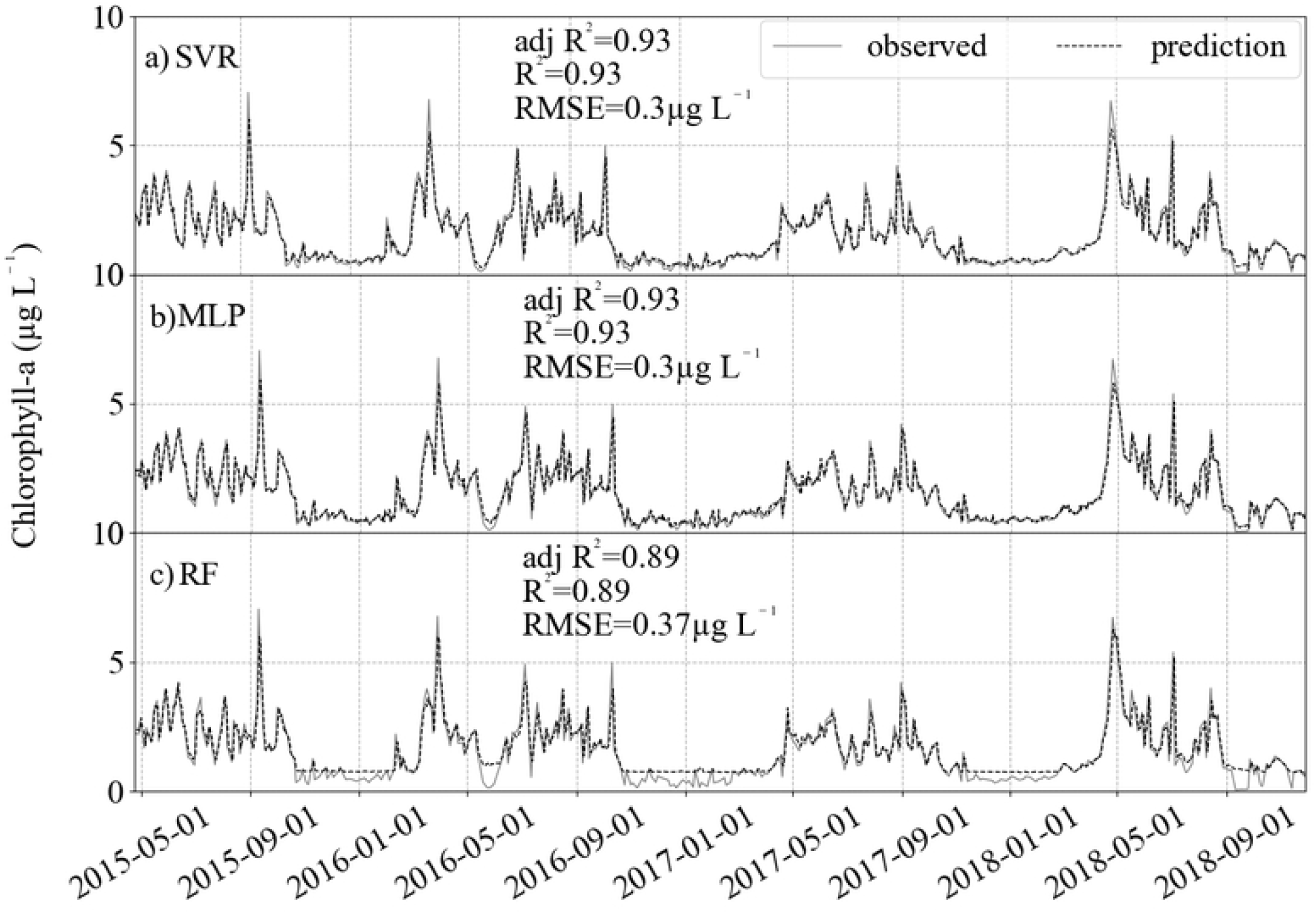
Result of prediction (black dashed) and comparison with the validation dataset (gray solid). For the three algorithms, R^2^ is approximately 0.9 and RMSE lower than 0.3 μg L^−1^. a) SVR, b) MLP and c) RF.

The iterative SARIMA parameters selection uses much more computer processing time compared with the GridSearchCV method in machine learning. The latter is a scale of seconds to minutes while the former hours to days. It took around two weeks to select the best p, q, P and Q parameters in the daily data considering a yearly seasonality. The results fitting the test dataset with the SARIMA model gave worst results compared with the ML models (Fig 7).

**Fig 7.**
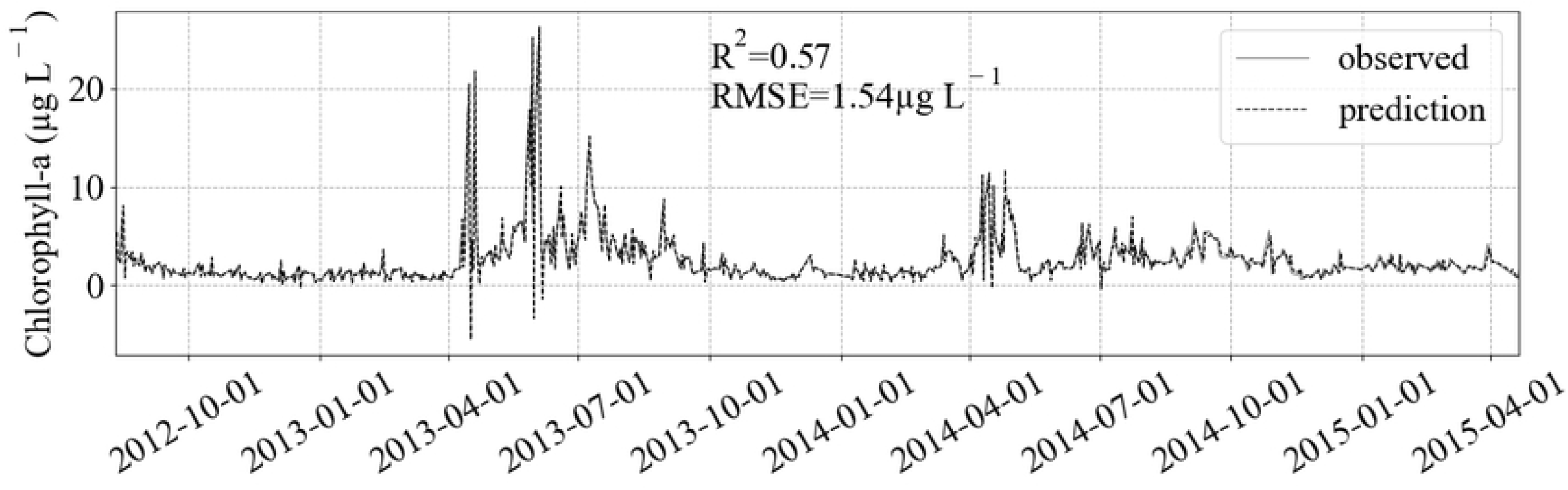
Result of SARIMA fit (black dashed) in the test dataset (gray solid). The better fit in extreme values is counter-balanced by the estimation of negative values, decreasing/increasing R^2^/RMSE compared to the ML models results.

## Discussion

Machine learning analysis was conducted on the Helgoland Roads Time Series to develop the best fit of chlorophyll-a concentrations over time using different parameters and their lagged correlates. For the three algorithms implemented, the model results were virtually equal in the evaluation metrics, presenting similar results in prediction, with slightly better values for the model SVR. For the time predictions, all the three models performances are acceptable with high R^2^ values greater than 0.70 and RMSE lower than 1.5 μg L^−1^, ~40% smaller than the chlorophyll-a concentration standard deviation of 2.9 μg L^−1^. However, all the algorithms were unable to predict extreme values (Fig 8). It was expected that a certain degree of decrease in accuracy would be incurred because of the difficulty in capturing and reproducing these extreme peaks [42]. One hypothesis which would explain the underestimation of extreme values is the lack of parameters evaluating e.g., hydrodynamics can result in the transport of chlorophyll from other areas as an input event, even though salinity and wind parameters are reliable indicatives for current and wave dynamics in the German Bight [43]. As these events do not present as a temporal pattern, the ML models does not recognize the influence in the target.

**Fig 8.**
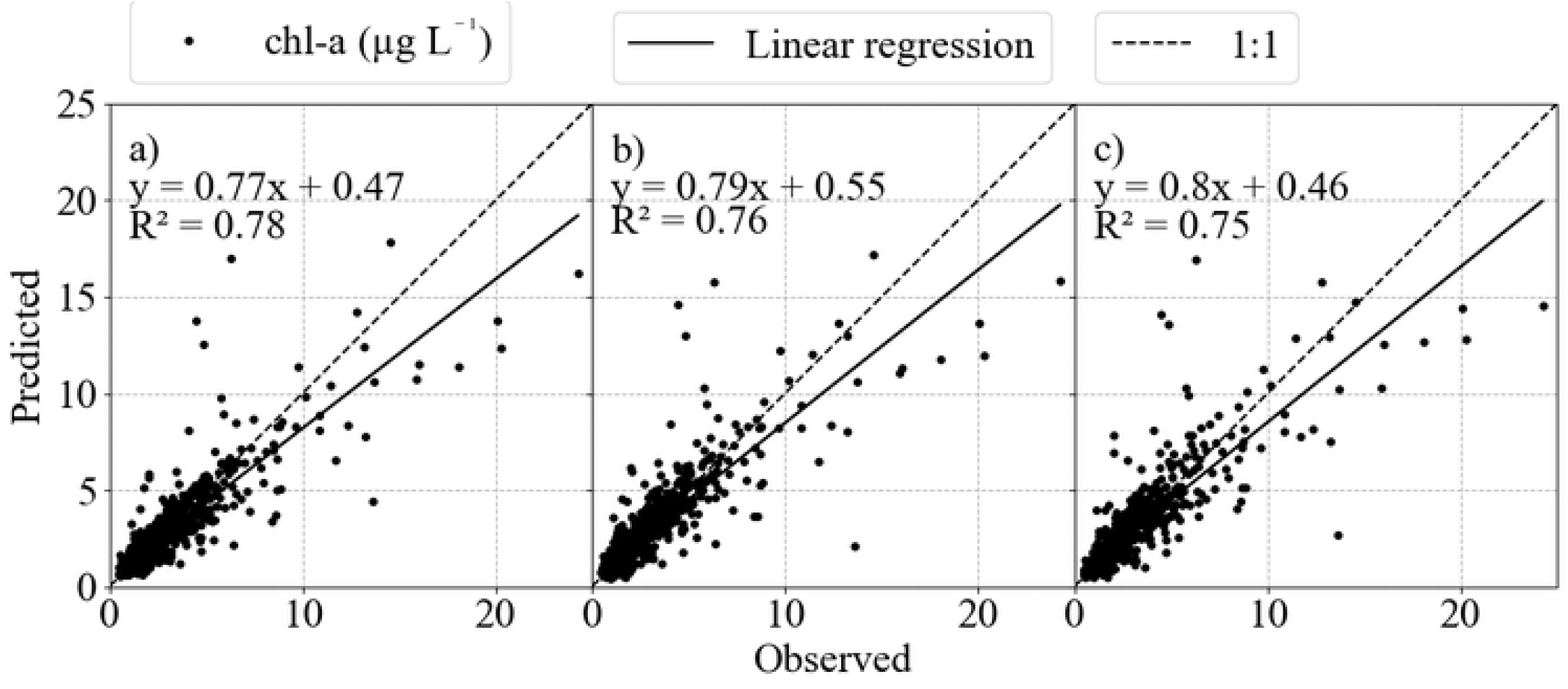
Cross-plots of the modeled and observed chlorophyll values in a) SVR, b) MLP and c) RF. It is possible to notice the deviation in extreme values, showing the limitation of the ML models in deal with these data values.

Because each algorithm is based on different algebraic assumptions and procedures, they can result in different predictions. Between SVR and MLP, [14] point to differences in the nonlinear equalization performance and the structural risk minimization principle of SVR being more effective than the empirical risk minimization principle of neural networks in terms of minimizing error. According to [44], in MLP the method for determining global solutions is difficult to converge because of its inherent algorithm design and model parameters are more complex than SVR, whereas the SVR has ready access to global optimal solutions, obtained by solving a linearly constrained quadratic programming problem [14]. Between SVR and RF, as we saw, the linear base model gave good results. There is the possibility of a linear dependency that is better captured by SVR, probably a result from the linear interpolation in the pre-processing step of this study.

The feature selection and tuning of hyperparameters was extremely important and improved the results substantially. This was noticeable in the adj R^2^ results for Default and Optimized models. While Random Forest gave the best R^2^ and RMSE using all features plus 2 lags, when fed with the 17 features selected by Ridge and SVR RFE, the estimation of extreme peaks was slightly better, but resulting in worse R^2^ and RMSE. It is the trade-off between variance and bias well discussed in the ML field. Analyzing the 17 features used in SVR and described in the Results section, the algorithm considered SST, lagged SST, lagged chlorophyll, Salinity and Secchi depth to reach the best results presented in this work. It is important to point that ML is a data-driven approach, but it is possible to make inferences about the selected features. For this study, we noticed the choice of SST as important feature, probably representing the seasonal patterns in the chlorophyll target.

Better R^2^, adj R^2^ and RMSE results in the independent validation dataset are possibly due to less variability and absence of extreme values, and shows the good generalization the ML models are capable. All the good results, for both the test and independent validation data, shows the better prediction power of the three ML algorithms evaluated in this study. Comparing with the classical SARIMA model, the univariate and linear background did not achieve results needed for it to outperform the ML models. Compared with the ML literature, studies like [3] and [11] achieved results of R^2^ ranging from 0.50 to 0.80, analyzing shorter time series of chl-a in lakes. [45] predicted variations of chlorophyll-a in different sites of the North Sea using Generalized Additive Models (GAM) and the R^2^ results ranged from 0.15 to 0.63. [28], using GAM to predict chl-a in a spatial approach for the North Atlantic, got the best result for R^2^ as 0.83. All these values show how variable can be different methods performances in predicting chlorophyll, not necessarily meaning one method is better than the other, but more adaptive. ML models proved generalization capacity and high accuracy.

## Conclusions

In this work, we evaluated three machine learning algorithms in a regression task. Support Vector Regressor presented a slightly better performance, with the advantage that it uses less computational time, and generated chlorophyll concentration predictions with 0.78 correlation to the observed data, in comparison to 0.77 and 0.76 to MLP and RF, respectively. Moreover, the root mean square error was around 1.1 μg L^−1^ for the test dataset and less than 1 for the independent validation data, which is approximately 38% percent smaller than the standard deviation of 2.9 μg L^−1^. This study demonstrates the ability of machine learning models to use environmental in situ time series to predict the chlorophyll concentration with significant accuracy (R^2^), higher than 70%, besides the importance of tuning hyperparameters and define the best predictors. Most chlorophyll-a prediction studies are conducted in fresh water environments or using satellite data and limited time series, so this work can be considered a step toward the use of Machine Learning algorithms in marine areas based on long term time series. Being aware of limitations presented in this study, in future works it would be interesting to work with irregular sampled time series, improve the method for feature selection and ensemble results of different ML and classical statistical models.

## Acknowledgements

We acknowledge the present and past crews of the research vessels Aade and Ellenbogen of the Biologische Anstalt Helgoland (BAH) for their unfailing provision of samples. We also thank Kristine Carstens, Silvia Peters, Ursula Ecker and all colleagues of time series group and those who were instrumental in the analysis of samples during the last decades. We thank the NOAA Physical Sciences Laboratory and Deutsche Wetterdienst for providing climate and meteorological data. This study was funded by the AWI Section Coastal Ecology.

## Supporting information

**S1 Fig**. Time series of parameters used to predict chlorophyll-a concentration: (a) Secchi disk depth, in meters (m); (b) Sea Surface Temperature, in degrees Celsius (°C); (c) Salinity; (d) Silicate (μmol L-1); (e) Phosphate (μmol L-1); (f) Nitrate (μmol L-1); (g) Sunlight duration, in hours (h); (h) NAO index; (i) Wind Direction, in degrees (°); (j) Wind Speed, in meters per second (m s-1); and (l) Total zooplankton abundance, individuals per cubic meter (# m-3).

